# Evolution of objectives can enable prosocial behaviour without social awareness

**DOI:** 10.1101/2025.08.06.668887

**Authors:** Bernd Meyer, Yue Yang

## Abstract

Division of labour is fundamental to the functioning of societies and socially living organisms. While it has been central to their study for decades, no complete picture has emerged yet. Some of the most fascinating questions arise in the context of self-organised societies, like those of social insects, that coordinate their behaviour with completely decentralised simple decision-making performed by individuals that only have local information at their disposal. Based on empirical evidence, these collectives appear to balance task engagement globally across their whole task network for the benefit of the colony overall. How can this pro-social coordination be achieved by independently acting individuals? How is a global workforce balancing possible based on only local perception with no knowledge of the global colony status or needs? Central to solving these problems is the question how can the relevant information flow through the task network so that a changed task demand in one part of the colony can lead to adjustments in distant other parts? We detail a model that presents a potential answer to this conundrum. Our model is informed by evolutionary game theory and rests on the assumption that the perception of an individual’s sensory input can evolve. We present simulation studies and a mathematical proof to show that pro-social behaviour will evolve in a collective of agents that adjust their behaviour using primitive and biologically plausible learning mechanisms if we assume an evolving perception function.

## 1 Introduction

Division of labour is fundamental to the functioning of social organisms and has been central to their study for decades [1]. The separation of tasks among different individuals or groups within a collective allows for the efficient use of resources and increases the chances of survival for the collective as a whole [2, 3, 4]. Empirical studies have shown that division of labour is prevalent in many socially living organisms, such as ants, bees, termites, and even some mammals [5, 6, 7, 8, 9].

Colonies of social insect are one of the most important model systems for the study of collective behaviour in general, and division of labour in particular. They are well known for their intricate colony organization and their ability to handle a wide range of tasks simultaneously, including foraging, colony defence, nest construction, temperature regulation, and caring for offspring [6, 10]. The colony’s ability to effectively allocate its workforce to these different competing tasks, adapting flexibly to changes in both external conditions and internal needs, is often cited as a key to their astonishing ecological success [11, 12, 13, 14]. Understanding the underlying mechanisms that enable this flexibility is thus fundamental to understanding these social organizations and the functioning of complex social systems in general.

It is well established that both developmental and genetic factors significantly influence the division of labour in social insects [15, 16, 17]. Additionally, studies have shown that faster and self-organized mechanisms for division of labour exist within colonies, enabling them to rapidly adapt to shifts in task requirements [18, 19, 20]. We adopt the term *task allocation* for this fast process to distinguish it from the more general phenomenon of division of labour.^1^ It is well established that the process of task allocation generally responds to altered environmental conditions such as food abundance, predation threats, weather and climate conditions [21, 22]. Empirical research has also emphasized the significance of social context and interactions in shaping the task preferences of individuals [23, 24, 25]. The fast dynamics of task allocation is thus driven by the interplay between workforce distribution, interactions structure, and environmental factors [26, 27, 28, 29, 30].

As this collective plasticity is ultimately based on flexible individual behaviour, understanding the rules that govern the task choice behaviour on the individual level is core to deciphering the collective phenomenon.

Much of the research on task choice behaviour has focussed on the performance of a single task or a binary choice between a pair of tasks. However, the reality is far more complex: the colony is faced with a complex network of tasks, spanning foraging, brood care, defence, nest maintenance, thermoregulation, and many others. Fitness pressure dictates that the task execution has to be balanced globally across the whole task network for the benefit of the whole colony rather than just locally for the immediate benefit of a particular task as perceived from the perspective of an individual. Apart from the obvious evolutionary fitness argument for network-wide or colony-wide optimisation, empirical evidence of such network-wide balancing in real colonies exists [31] and has been known for almost 50 years [32, 33, 34] (summarised in [35]).

Arguably, the most puzzling aspects of global task allocation is that it must be managed by individuals that generally lack knowledge of the overall state of the colony,^2^ so that their behavioural decisions must rely on only local information that can be readily sensed by them [37, 38], potentially in combination with additional information obtained in interactions with other colony members [39, 40].

Questions about this global balancing can be approached from two different perspectives. The first perspective, which drives our work, concerns the proximate behavioural mechanisms: *How* is such global balancing even possible based on only local perception? *How* can information propagate through a task network, so that a changed task demand in one part of the task network can lead to adjustments in distant other parts? The second perspective it that of evolutionary theory, which asks *why* altruistic behaviour of individuals arises in the first place, without investigating the concrete mechanisms.

It turns out that the answers to both questions are tightly linked: they lie in the evolution of perception. What is missing in the overall picture is an explicit link between individual behaviour and integrative colony fitness which is not visible at the level of the individual. An evolving “perception function”, which shapes how directly perceived sensory inputs are used in the process of behaviour adaptation, can establish this link. *The single assumption of an evolving perception function is sufficient to let pro-social behaviour emerge within a collective*.

To study this, we consider an abstract task allocation scenario in which a group of agents has to balance global task engagement across a network of tasks. Individuals can migrate between tasks along the edges of this network. At any point of time, an individual is only concerned with a binary choice between two tasks and can only sense information about this pair of tasks.

Such binary choices are a paradigmatic scenario, both in empirical work and in modelling studies. In our model, each agent in the collective independently decides which tasks to execute and adapts its task choice via a primitive learning mechanism by exploring a new task and continuing to engage in the one for which it perceives better task feedback on the individual level.

We first investigate a binary choice scenario and show how the agents learn to solve simple allocation problems using only locally perceived information but inevitably fail to act pro-socially (in the interest of the collective) in more difficult scenarios. We then introduce an evolving “perception function”, modelled as an evolvable neural network, that shapes how the locally perceivable and directly sensed information is used as input to the learning process. Since this effectively amounts to the reshaping of the objective function of an optimisation problem (see Section 2), we term this process “evolution of objectives” or “objective evolution” for short.

For the binary choice scenario, it turns out that this simple and plausible addition enables the agents to learn pro-social behaviour, because it establishes a link between colony-level fitness pressures and individual behaviour.

Next, we show that the ability to “correctly solve” binary task choices also resolves the network-wide balancing problem as it enables task-demand information to propagate unhindered through the network. Interestingly, it turns out that the propagation of demand information only works fully if individuals do not always rigidly adopt the better tasks but sometimes stay with the task that yields inferior feedback which may happen due to less stringent individual decision making or noisy task feedback. Conceptually, this is closely related to the role of noise in the adaptive decision making of social insects in other, simpler cases [41].

Effectively, our model shows that we do not have to assume complex forms of information exchange between individuals, since global regulation can already be achieved on the basis of propagating task demands.^3^ Finally, we show the applicability of our model to real biological scenarios in the context of the transport-optimising behaviour of foraging Army ants.

It is important to note that we do not claim to have biological evidence that this is the actual process taking place in social insect colonies. However, this is a parsimonious, consistent, and, we claim, biologically plausible model of how global pro-social task balancing in a network of tasks can take place, a process for which currently no general model exists. We thus believe that it is a hypothesis worthy of further investigation. The point of the present paper is to detail this hypothesis and show through mathematical proof and computational simulations and that it is a possible explanation consistent with empirical knowledge.

## 2 Model

Our model captures how task allocation can be regulated to achieve pro-social outcomes even if rational individuals independently adjust their task choices using only locally perceived information. We construct this model in two steps: In the first step, the model addresses a single binary choice between two tasks, while the complete model addresses a network of such binary choices, so that the whole task network of the collective can be captured.

### 2.1 Base model for binary choice

#### 2.1.1 Individual agent behaviour

We consider a set of N_a_ agents 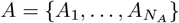 and a set of *N*_*t*_ tasks 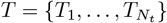 that these agents have to perform. Let 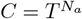 be the joint action space (task choices for all agents) and *Z* ∈ *C* be the joint action profile for all agents. We write [*Z*]_*j*_ to select the *j*-th component of a vector, ie. [*Z*]_*j*_ is the task that agent *A*_*j*_ engages in. As usual, we write *Z*_−*j*_ for the action profile of all agents other than *A*_*j*_ with *Z* = ([*Z*]_*j*_, *Z*_−*j*_).

Each agent *A*_*i*_ decides independently which task to engage in based on a behaviour policy 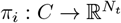 that maps the current task profile to the probability of engaging in any of the tasks currently accessible to *A*_*i*_.

Agents individually perceive task-specific performance feedback (or rewards), which could, for example, be the quality and/or the amount of food retrieved during foraging. We allow for the possibility of this feedback to be dependent on the task choices of other agents and specifically to be frequency-dependent.

The task feedback function 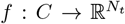 with 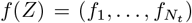 gives the feedback for all tasks, which are specific to each task but the same for all agents engaging in the same task. Ultimately, we are interested in the social benefit (the benefit for the whole collective) that we take as a fitness proxy. This is denoted by *F* : *C* → ℝ.

Central to our model is the fact that individual agents cannot directly perceive the social benefit *F*(*Z*). Individuals can only directly observe *N*_*S*_ sensory inputs. We denote the sensory input of agent *A*_*i*_ as 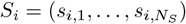 with *s*_*i,j*_ ∈ ℝ. Depending on the scenario modelled, the immediate task feedback, specifically for the task [Z]_i_ that the agent is currently engaged in, would typically be included in S_i_.

Crucially, we assume that these sensory inputs are not directly used to control behaviour adaptation but instead that they are first processed by a *perception function N*_*θ*_ : 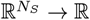 that we implement as a neural network with parameters *θ. N*_*θ*_ is thus an universal function approximator.^4^ In our model this function approximator evolves (see Sec. 2.1.2).

*It will turn out that, provided* S *is sufficiently informative, the ability of* θ *to evolve is sufficient for the collective of agents to evolve pro-social behaviour*.

As a shorthand, let *Ŝ*_*i*_ = *N*_*θ*_(*S*_*i*_) ∈ [0, 1] be the perception of agent *A*_*i*_, ie. its sensory inputs processed through the perception function and normalised in the unit interval.

Our model progresses in discrete time steps t_1_, …, t_n_.^5^ At each time step, all agents adjust their task choices based on their perception of task feedback with a very simple response behaviour, in which they seek to improve this feedback. We assume that they occasionally probe other tasks and switch to a new task with a high probability if it leads to better perceived feedback. However, with some baseline probability they remain in the original task regardless of feedback.

We can formalise this as *better reply with inertia* behaviour. Let *B*_*i*_(*Z*) be the better response set for agent A_i_, i.e. the set of alternative tasks that lead to higher perceived feedback.

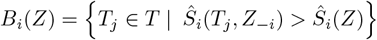

At each time step *t* each agent *A*_*i*_ chooses a task to engage in at time *t* + 1 according to *π*_*i*_(*Z*) defined as:

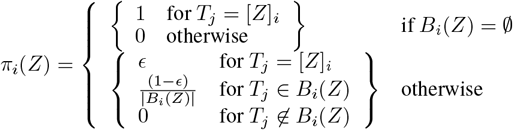

where ϵ ∈ (0, 1) is the agents’ inertia. We note that the fine details of the behaviour revision protocol do not matter as the asymptotic dynamics of learning in the class of games we are concerned with are inherently robust for all kinds of better reply processes as long as they use only finite memory [42].

#### 2.1.2 Colony evolution

The model so far captures the fast behaviour adaptation by individuals on short time scales during their lifetime. Crucially, the parameters of the perception function can evolve. The dynamics of the system thus plays out on two different timescales. On the fast time scale (behaviour of the collective), we assume *θ* as fixed and let the agents’ behaviour equilibrate resulting in the joint behaviour policy 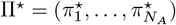. On the slower timescale (evolution of the collective), we assume 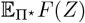 as the fitness at steady-state and let θ evolve accordingly through mutation and crossover using differential evolution [43].

#### 2.1.3 Environment (binary choice)

The task of pro-social reward optimisation is trivial if independent individual rationality leads to the pro-social optimum. We are specifically interested in cases where the benefit expectation in equilibrium under individual rational choice is less than the pro-social optimum, ie.

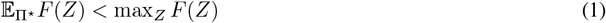

Congestion games, a well-established model of *n*-player social dilemmas [44] in which Nash equilibria are not pro-social, are a suitable model of this. We thus define our environment first as a single congestion game and later as a network of congestion games (see Section 2.2). A congestion game generally models a situation in which players make use of a set of capacity-limited resources. Each action that a player can choose requires the use of one or more resources and there is a penalty for resource use. This penalty increases with the number of players utilising the same resource. In other words, the payoff for a player is inverse in the number of players utilising the same resource(s).

This is, for example, exemplified in foraging and transport scenarios where overcrowding can reduce the efficiency of individuals [45, 46].^6^ More generally and more importantly, we can also take this as an abstract model of a task choice in which two tasks have to be balanced rather than individually optimised, for example different food sources due to nutritional geometry [48].

In our setting, each individual congestion game represents a binary choice between two tasks (for example foraging at one of two sources, N_T_ = 2). We call this an *atomic* game. The individual reward for each task is frequency-dependent and monotonically decreasing in the number of agents engaging in this task. The specific choice of the reward function is not crucial. For the purpose of our experiments and motivated by the fact that the logistic function 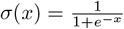 is a suitable fundamental choice for a capacity limited resources, such as the yield of a food patch [49], we define

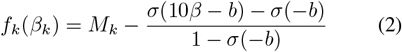

where *β*_*k*_ is the proportion of the agent population engaging in task *T*_*k*_, *M*_*k*_ is the maximum individual payoff that can be obtained from task *T*_*k*_ and b is a parameter determining the inflection point of *f*. With this, the payoff for the entire group of agents becomes

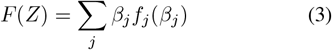

where *β*_*j*_ is the population fraction engaging in task *T*_*j*_ under the joint task profile *Z*. See Figure 2 for the relationship between these functions. Learning mechanisms in which individuals try to optimise their own feedback, such as the one given above, will drive the system to the Nash equilibrium, which is clearly not the social optimum.

Note that we have restricted atomic games to binary choices only for simplicity and to save notation. The extension to *N*_*T*_ > 2 is straight forward.

### 2.2 Model for task networks

So far we have considered binary choices. While this could be extended to ternary or higher-order choices in a straight-forward manner, it makes more sense to consider a network of tasks for the general case. From a modelling perspective, this corresponds naturally to agents having a restricted set of local task-choices that changes as they are moving between tasks.

We assume there is a reachability relation between tasks (ie. agents can migrate between some tasks but not others) modelled as an undirected graph 𝒢 = (*T, E*) with tasks as vertices as in Figure 1.

**Figure 1:**
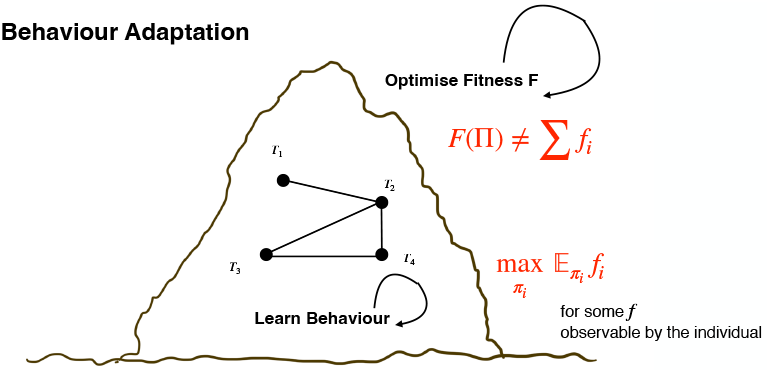
In the classical model, behaviour adaption amounts to individuals learning behavioural policies π_i_ that optimise the expectation of their individual task reward (or performance feedback) f_i_, for example, the amount or quality of food retrieved in foraging. However, there is no channel for information flow from the overall colony fitness F, which depends on the joint behaviour Π of all agents, to the individual feedback. If F is not simply additive across all tasks and individuals, this can lead to behaviour policies that are not pro-social.

**Figure 2:**
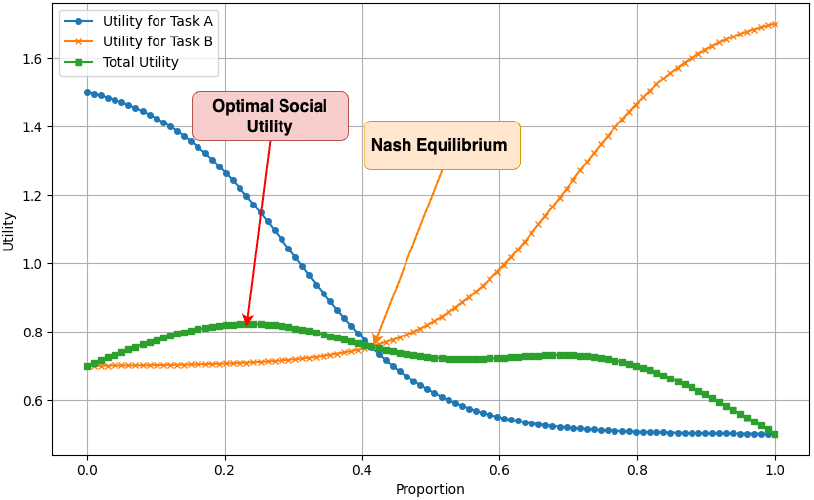
Individual and collective payoff for a binary task choice: The maximum of the collective payoff (green curve) does not coincide with the Nash equilibrium (intersection of the individual payoffs). Thus, the Nash equilibrium is not pro-social.

At each point in time an agent participates in one particular two action game G_i_ that models a choice between two tasks. Agents can migrate from task *T*_*i*_ to *T*_*j*_ if there is a corresponding edge (*T*_*i*_, *T*_*j*_) ∈ *E*. We assume that 𝒢 is connected, so that agents can ultimately reach any task in the network. Note that each task can be a simple task or an atomic game *G*_*i*_ as defined in Section 2.1. In the latter case we assume a separation of time scales for the convergence of the atomic game and the network game.

𝒢 induces a state-based game [50], a simpler form of Markov games [44]. It is defined as *M* = (*A, C, U, X, P*), where A is the set of agents, *C* the joint action space,^7^ 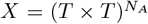 the state space, 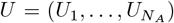 with *U*_*i*_ : *C × X* → ℝ the state-dependent utility function for all agents, and P the state transition kernel.

The state space X captures in which game each agent currently participates. We initially investigate transition dynamics in which an agent moves through the task network, actively searching for a better task. This corresponds to games in which it continues to move to new tasks and compares this to its original tasks until it can switch to a better one. Importantly, we will relax this assumption in Sec. 4, since such behaviour may not be realistic in large task networks.

Let *G*^*+*^ define the set of games reachable under action a from a given game:^8^

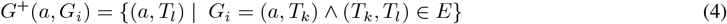

The state transition probability *p*_*s*_ : *X × T × X* → ℝ captures how agents migrate between games. The only necessary requirement is *p*_*s*_(*G*_*i*_, *a, G*_*j*_) > 0 iff *G*_*j*_ *∈ G*^+^(*a, G*_*i*_). We may, for example, simply assume a uniform distribution.

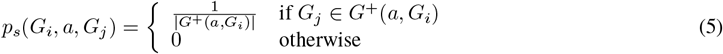

The transition kernel *P* : *X × C × X* → [0, 1] for the whole game allows a single agent to move at a time.^9^

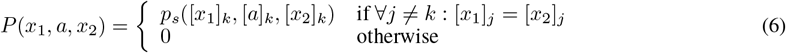

Let #_*i*_(*a*) be a function that counts the agents performing task *T*_*i*_ in joint action *a*. The utility of agent *A*_*k*_ in state *x* is defined as the total utility of the tasks in the atomic game that it currently participates in.

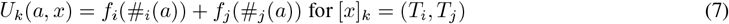

We initially assume that the one-step utility of the entire network game is the sum of the utilities of all tasks in the network. However, other forms of combining individual task utilities are possible. Of particular biological relevance is a combination in which some tasks are limiting the overall benefit. For example, we may want to capture the situation that no task can ultimately generate a fitness benefit if brood care is entirely neglected. Such situations can be modelled by a *minimum* instead of an additive utility, which leads to a *maximin* optimisation problem instead of a standard maximisation problem. This can be handled in similar forms. We leave the details for future work.

## 3 Simulation Results

We first simulate binary choice between two congestive tasks in an atomic game as described in Section 2.1.3 and then proceed to simulate complete task networks.

### 3.1 Binary choices (atomic games)

For binary choice scenarios, we use the binary congestion problem (N_T_ = 2) defined in Section 2.1.3 with ten randomly drawn parameter sets. These parameter sets are given in Table 1. We use multiple problems to show that a single unified perception function can be evolved that applies to different settings rather than just one particular scenario.^10^ We then apply the evolutionary procedure defined in Section 2.1.2 to colonies in which individuals behave as defined in Section 2.1.1.

**Table 1:** Parameter list for benefits and costs for A and B.

| Task | Benefit ($B_A, B_B$) | Cost for A ($C_A$) | Cost for B ($C_B$) |
| --- | --- | --- | --- |
| 1 | 2 | 0.3 | 0.5 |
| 2 | 2 | 0.5 | 0.3 |
| 3 | 2 | 0.6 | 0.2 |
| 4 | 2 | 0.4 | 0.2 |
| 5 | 2 | 0.8 | 0.6 |
| 6 | 2 | 0.1 | 0.3 |
| 7 | 2 | 0.9 | 0.7 |
| 8 | 2 | 0.5 | 0.6 |
| 9 | 2 | 0.7 | 0.9 |
| 10 | 2 | 0.2 | 0.6 |

We evolve a population of perception functions, the fitness of which is defined as the difference between the maximally achievable collective payoff and the collective payoff achieved in the steady state by a population of agents adhering to better-reply dynamics based on this perception function.

Note that the perception function is only used to drive the learning dynamics and that the actual payoffs entering into the fitness calculation are derived from the true collective payoff as follows: Let *β*^⋆^ = argmax_*β*_ *F*(*β*) be the population fraction for the first task that maximises the collective fitness.^11^

Let 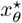 be the steady-state proportion of the population engaging in the first task as it results from better-reply dynamics according to the perception function *N*_*θ*_. The fitness *H* of this perception function is then defined as:

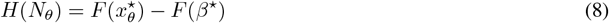

The resulting perception functions for the parameters in Table 1 are compared with the true objectives in Figure 3. We observe that in all cases, a (almost) unimodal perception function evolves with an *argmax* that is is very close to the *argmax* of the true objective, inline with our hypothesis.

**Figure 3:**
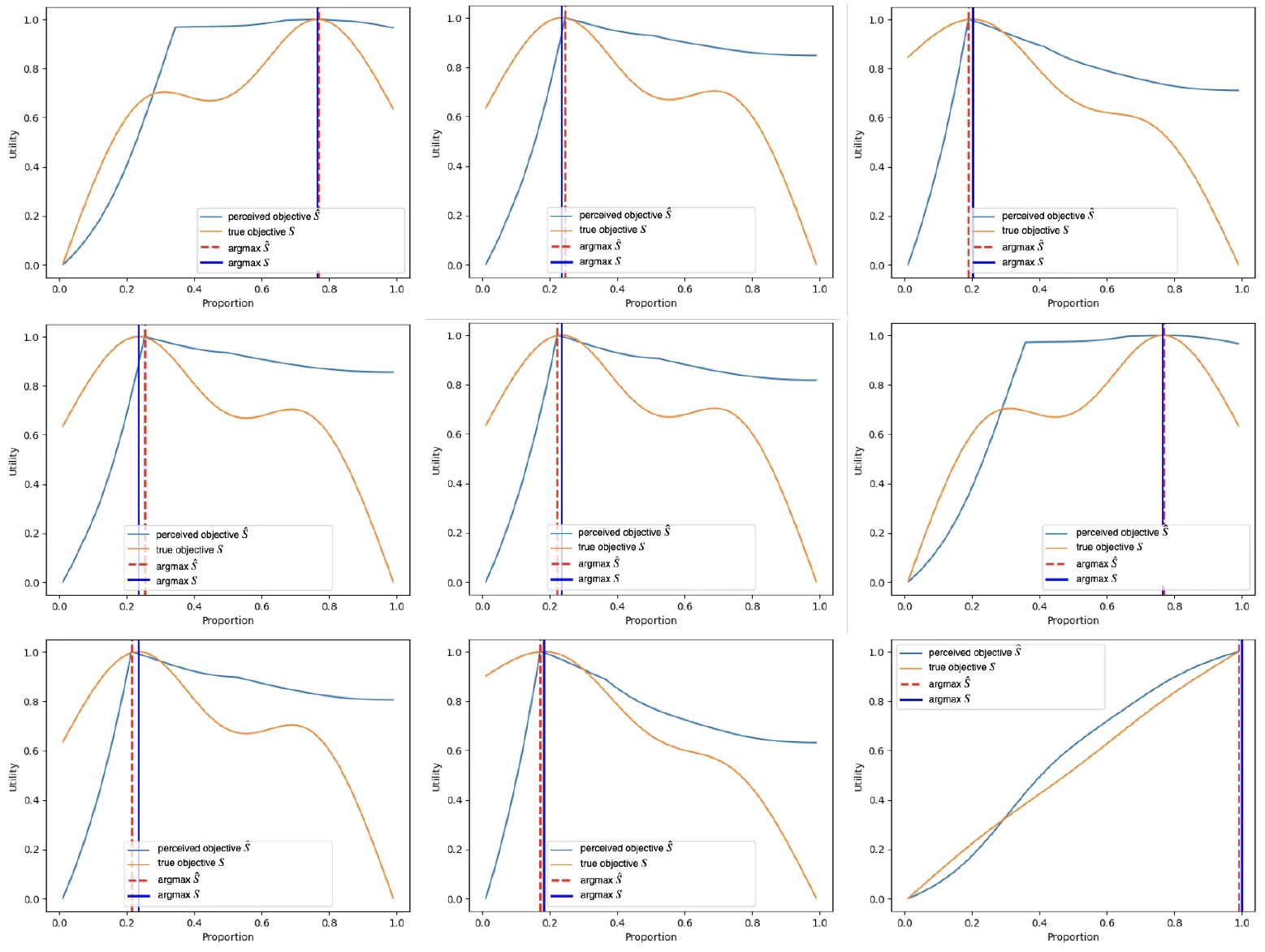
Comparison between the true collective objective and the perceived objective for different objective parameter settings: The perceived objective (blue line) generally evolves to be a unimodal function with the global optimum at a point that is close to the global optimum of the true objective (orange line).

Note that these simulation should really only be understood as a sanity check. It is easy to see abstractly that the outcomes cannot be different. There is selective pressure on the global *argmax* of the perception function to be the same as that of the true objective. This is because the true collective payoff is optimal under this condition, implying maximum fitness. Furthermore, there is selective pressure on the perception function being unimodal as the fitness is highest if the *argmax* is reached by stochastic gradient ascent from every possible starting point. Only a unimodal perception function allows hill-climbing to reach the optimum from any starting point. Thus, a unimodal perception function that has its optimum at the same point as the true objective has the optimal fitness and should normally evolve. Of course, there is no guarantee that an evolutionary process will find the global optimum. In our case, we might expect this to fail if the true objective has multiple local optima that are close in their objective values. In such cases, the stochastic nature of the perception function evolution might develop towards a local optimum.

Since all agents share the same utility function, better reply with inertia dynamics using this perception function will converge on the maximum of the perception and hence on the pro-social maximum. It is important to contrast this with the outcome that would result if we eliminated the perception function and let the agents perform better-reply hill climbing using the immediate task reward [*f*(*Z*)]_*j*_ instead to adapt their behaviour when performing task *T*_*j*_. We know that the steady state would be a Nash equilibrium. Independently of the specific parameters, all our reward functions have the shape shown in Figure 2. The behaviour adaptation would thus equilibrate where the payoff for both tasks is equal [*f*(*Z*)]_1_ = [*f*(*Z*)]_2_. As is evident in the figure, this is generally not the point of optimal collective payoff.

In summary, the collective of agents has evolved a perception function that, independently of the particular parameters of the environment, allows them to optimise the collective utility (fitness) for the colony without requiring explicit coordination or even explicit awareness of the social context. This optimum would not be reached if the agents exhibited the same behaviour but used the immediately sensed utility of the task executed. From the technical perspective of game theory, the perception function has reshaped the atomic game into a potential game [50] with a Nash equilibrium that coincides with pro-social behaviour.

Thus, simply introducing an evolvable perception function allows the collective to evolve pro-social behaviour without requiring any additional social coordination mechanisms beyond the mechanisms already used for the control of solitary behaviour.

### 3.2 Task networks

We now show that we can extend the approach from interactions that are limited to a single (fully connected) task group to a network of tasks as defined in Section 2.2.

Our simulation network comprises *N*_*T*_ = 8 tasks in *N*_*G*_ = 4 atomic games. This is because four atomic games is the smallest configuration in which an investigation can be performed across different topologies. We investigated 600 scenarios, corresponding to 100 random parameter sets for each of the possible six topologies (see Fig. 4). We use *αe*^−*βx*^ with 0 < *α, β* ≤ 1 as the congestion function. All simulations run for 25,000 update steps. We established a performance baseline by determining the network payoff for 10,000 random parameter sets for each configuration.

**Figure 4:**
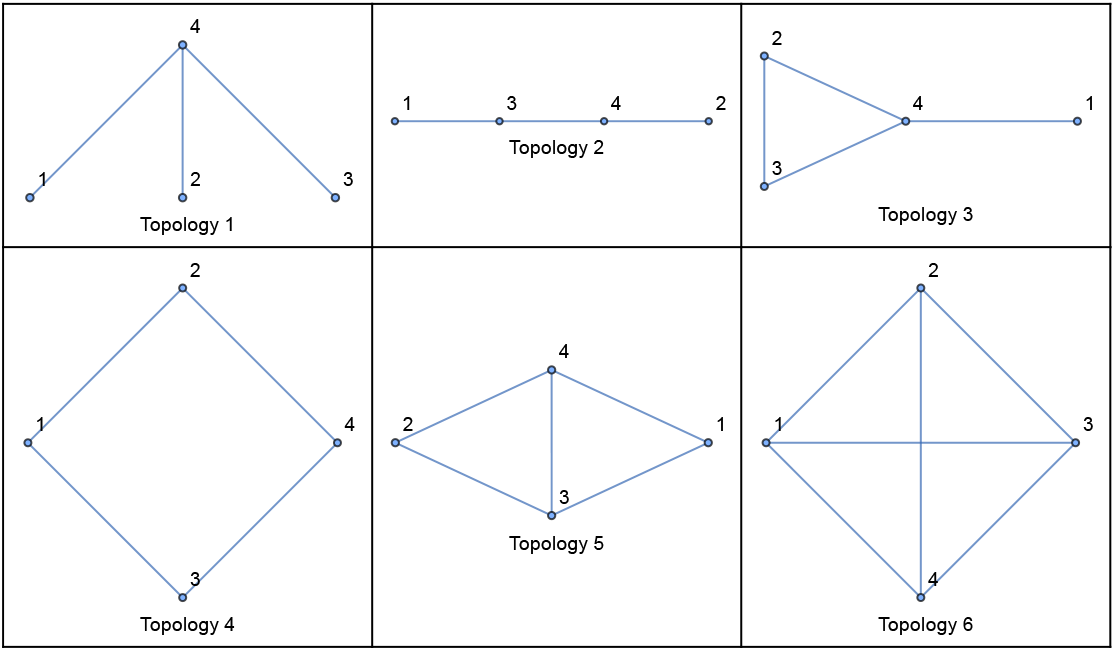
Game network topologies. Vertex labels correspond to the indexes of atomic games in the example runs detailed in Fig. 7.

Optimality in the table and all figures is measured as the fraction achieved of the theoretically possible payoff improvement over the minimal payoff for these parameters:

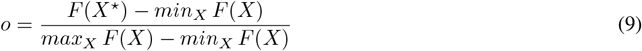

All runs exhibit the same network payoff dynamics that is increasing over time and concave but they converge at different levels of optimality (Figure 5). Obviously, in the fully stochastic setting not every simulation can reach the optimum. However, the final payoff is significantly above the baseline in the vast majority of cases (Table 2) and we rarely see a process stuck at low payoffs (Figure 6). No significant differences between different topologies can be observed.

**Table 2:** Comparison of optimality to baseline across all topologies.

| Topology | Optimality Mean (n=100) | Optimality Std Dev | Baseline Mean (n=10000) | Baseline Std Dev |
| --- | --- | --- | --- | --- |
| 1 | 0.875 | 0.093 | 0.599 | 0.118 |
| 2 | 0.897 | 0.086 | 0.598 | 0.118 |
| 3 | 0.902 | 0.082 | 0.600 | 0.118 |
| 4 | 0.887 | 0.090 | 0.599 | 0.118 |
| 5 | 0.909 | 0.078 | 0.599 | 0.119 |
| 6 | 0.901 | 0.078 | 0.598 | 0.118 |
| **All** | **0.895** | **0.085** | **0.599** | **0.118** |

**Table 3:** Key measurements.

| Symbol | Definition |
| --- | --- |
| $N$ | Total number of ants available on the foraging trail. |
| $L_T$ | Length of the main trail (not including the experimental gap). |
| $L_E$ | The length of the inner edge of one arm. |
| $L_1$ | The Length of apparatus arm that overlaps with the other arm. |
| $L_0$ | The length of the arm between the two hinge points. |
| $L_A$ | Effective inner length of the apparatus (the “gap”), which depends on the angle $\theta$. |
| $d$ | Distance the living bridge has moved from the junction of the bridge toward the main trail axis. |
| $D_{\max}$ | Maximum possible distance $d$ can reach (i.e., the full shortcut). |
| $b = 2d \tan(\frac{\theta}{2})$ | The physical length of the bridge once it has extended a distance $d$. |
| $w_\theta$ | Empirically measured ratio of bridge width to bridge length (depends on $\theta$). |
| $\ell_n, w_n$ | Length and width of an individual ant (used to estimate how many ants form the bridge). |

**Figure 5:**
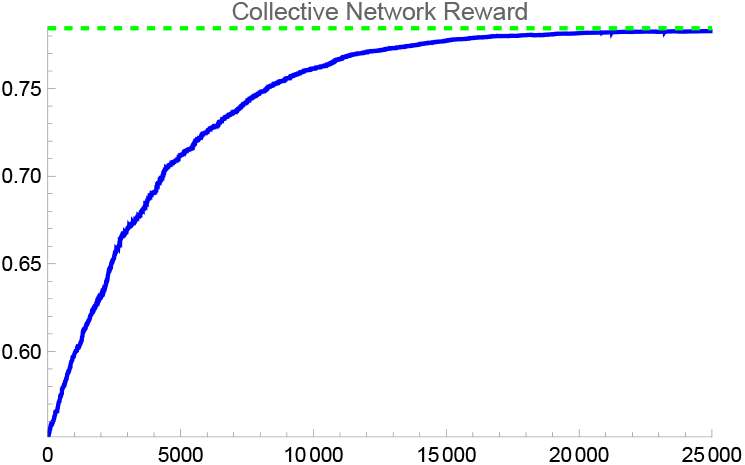
Network payoff over 25,000 update steps. The vertical scale is the absolute network payoff (not the optimality level) and the green line indicates the theoretically achievable maximal payoff.

**Figure 6:**
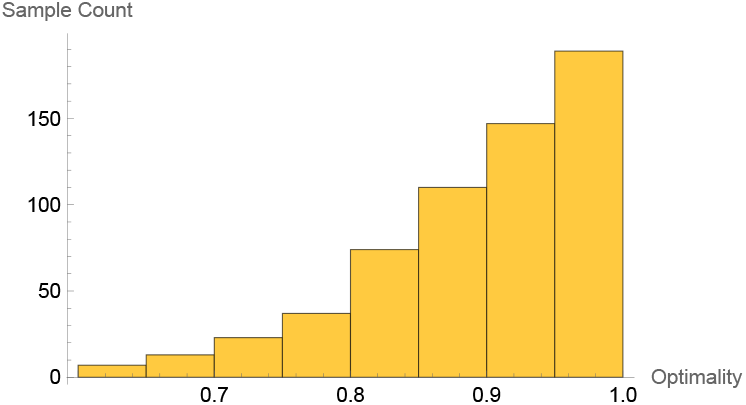
Simulation outcomes for 600 runs with 25,000 update steps each across all 6 topologies.

We give examples of two typical runs for each topology, one converging close to the optimal distribution and one converging to a different distribution (Figure 7). Upon closer examination, at least one of the following two causes seems to be involved in almost all cases where the network converges to a distribution that is significantly different from the optimal one:

**Figure 7:**
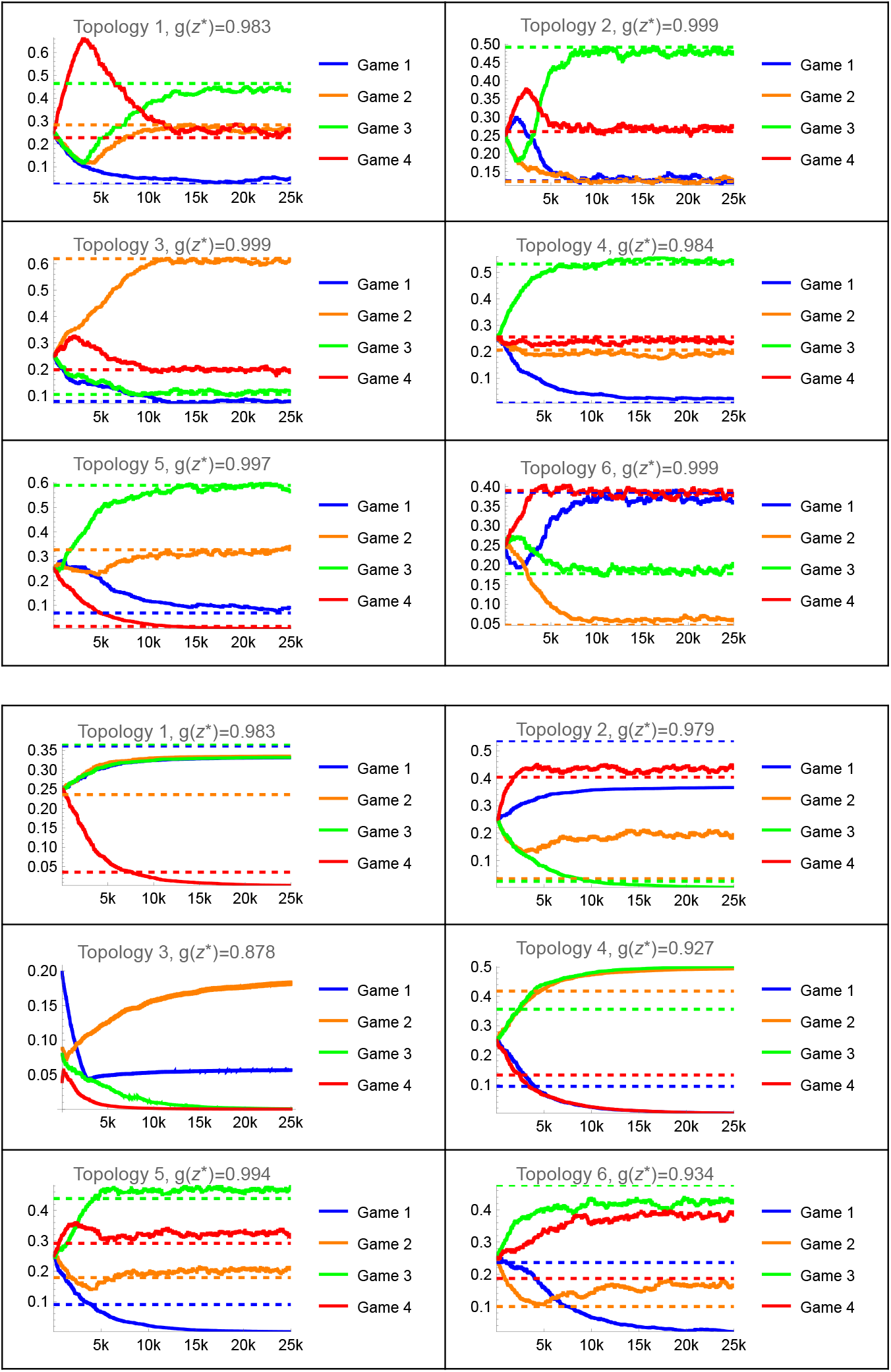
Distribution of the agent population across games over time for sample runs that converge on the optimal distribution (top panel) and on suboptimal solutions (bottom panel). Solid lines indicate the fraction of the population in the respective game, dashed lines indicate the fraction at the theoretical optimum of the problem for the given parameters. Game indices correspond to the vertex labels in Fig. 4.

- *Node crash*: When the occupancy at a node becomes very low, the improvement in objective value that agents moving from this node to neighbouring nodes can achieve may not be large enough to overcome the random fluctuations of the stochastic process on the timescale of the simulation. This can be particularly pronounced at articulation points that can disconnect the graph (Examples: Topology 1–3).
- *Lack of pressure*: When the potential landscape is relatively flat, the gradient may not be large enough to overcome the stochastic fluctuations on the timescale of the simulation. In these cases the objective value is often still close to the optimum but the distribution of agents across games is different. Examples: Topologies 1, 2, 4–6.

## 4 Convergence in Task Networks

Next, we prove that better response with inertia dynamics stochastically optimises the social benefit in the task network game. The convergence argument rests on the fact that M is a state-based potential game [50].

We have defined *M* as a state-based game in Section 2.2. To show that *M* is a state-based *potential* game (Def. 3.2 [50]), we need to ensure that there exists a potential function Φ : *C* × *X* → ℝ such that for every action-state pair (*a, x*) ∈ *C* × *X*:

i. For any agent *A*_*i*_ and action 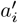: 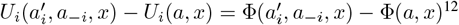^12^
ii. For any state x^*′*^: *P*(*x, a, x*^*′*^) > 0 ⇒ Φ(*a, x*^*′*^) ≥ Φ(*a, x*)

The first condition is the usual potential condition adapted to state-based games. The second condition requires all action invariant state trajectories to have non-decreasing potential.

Let the potential function be

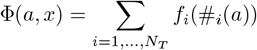

The utility of agent *A*_*i*_ in state *x* with [*x*]_*i*_ = (*T*_*k*_, *T*_*l*_) changes with a unilateral deviation from [*a*]_*i*_ to *a*^*′*^ as

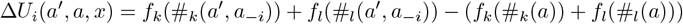

and *A*_*i*_ cannot change the payoff of any tasks other than *T*_*k*_, *T*_*l*_.

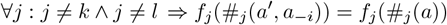

With

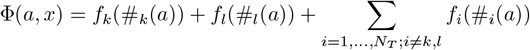

the change in potential with a unilateral deviation of *A*_*i*_ from [*a*]_*i*_ to *a*^*′*^ is

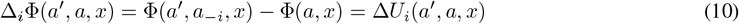

Hence, condition (i) is satisfied.

Condition (ii) is trivially satisfied since Φ is state independent by definition,^13^ i.e. ∀*x, x*^*′*^ ∈ *X* : Φ(*a, x*^*′*^) = Φ(*a, x*). *M* is thus a state-based potential game. Theorem 4.2 of [50] then guarantees that M will almost surely converge to an action-invariant set of recurrent state equilibria if all players adhere to finite memory better reply with inertia learning. What remains to be shown is that the recurrent state equilibria maximise the collective utility.

The definition of recurrent state equilibria is based on the concept of action invariant state trajectories. For a state action pair [*a*^0^, *x*^0^] the set of reachable states on an action invariant state trajectory is *X*(*a*^0^ | *x*^0^) ⊆ X where *X*(*a*^0^ | *x*^0^) = {*x* ∈ *X* | ∃*t* > 0 : *Pr*[*x*(*t*) = *a* | *x*(0) = *x*^0^] > 0} conditioned on the process *Pr*[*x*(*t* + 1)|*x*(*t*) = *x*] = *P*(*x, a, x*(*t* + 1)) for an action dependent state transition kernel *P*. Since the transition kernel of *M* is action independent *∀*a : *P*(*x, a, x*^*′*^) = *P*(*x, x*^*′*^).

A recurrent state equilibrium is a state action pair [*a*^⋆^, *x*^⋆^] for which

i. *x*^⋆^ never becomes unreachable on any action invariant state trajectory from [*a*^⋆^, *x*^⋆^], ie. ∀*x* ∈ *X*(*a*^⋆^ | *x*^⋆^) : *x*^⋆^ ∈ *X*(*a*^⋆^ | *x*)
ii. all agents play their best response to all other agents in all states *x* ∈ *X*(*a*^⋆^ | *x*^⋆^).

Condition (ii) implies that Φ(*a*^⋆^) is maximal. To show this, assume that [*a*^⋆^, *x*^⋆^] is a recurrent state equilibrium in which Φ(a) can be increased. By Eq. 10, there is an agent *A*_*i*_, an action *a*^*′*^ and a state x with ∀*j* : [*x*]_*j*_ = ([*a*]_*j*_,·) and [*x*]_*i*_ = ( ·, *a*^*′*^) in which *A*_*i*_ can increase its utility by unilaterally changing their action from [*a*]_*i*_ to *a*^*′*^. By Eqs. 4, 5 and since 𝒢 is connected, *x* ∈ *X*(*a*^⋆^ | *x*^⋆^). *A*_*i*_ is not playing its best response to *a*_−*i*_ in x. This contradicts condition (ii) of the assumption that [*A*^⋆^, *x*^⋆^] is a recurrent state equilibrium. Thus, the potential Φ(*a*^⋆^) cannot be increased if [*a*^⋆^, *x*^⋆^] is a recurrent state equilibrium.

Hence, better reply with inertia dynamics maximises the potential Φ, which defines the collective utility.

The above proof rests on the model’s assumption that an agent continues to compare the same task with new trial tasks as it moves through the task network to find a better task to switch to. The required comparisons may not be realistic in a large task network where comparisons may only be possible locally, ie. between tasks that are adjacent in 𝒢.

When only local comparisons are possible any dynamics in which an agent can strictly switch only to a better response may not lead to convergence on a maximised potential. This is because an agent may not be able to find a better task even if it exists because its movement is blocked by inferior tasks along all paths to this task. For comparisons restricted in this way, only a transition dynamics that allows agents to occasionally move to inferior tasks with a positive residual probability can guarantee convergence to a maximal potential state.

We modify our model to only allow local comparisons by redefining *G*^+^ so that agents only ever compare their current action to another action adjacent to it in 𝒢:

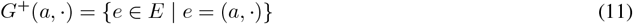

A common type of dynamics in which agents move to better actions with high probability and the probability to move to an inferior action decays rapidly with the difference in utility is log-linear learning [51]. It is defined in the following way: At each time step *t* at most a single agent *A*_*i*_ is chosen to explore a new task and there is a positive probability that no agent is chosen. A chosen agent *A*_*i*_ in state *x* picks an alternative trial action *a*^*′*^ uniformly from the alternative actions available to it and adopts the next task with the following probabilities:

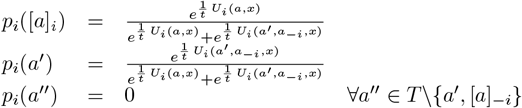

where *a* is the joint action at time *t* − 1.

Theorem 5.1 of [50] proves that under log-linear dynamics in an exact^14^ state-based potential game the state action pairs [*a*^⋆^, *x*^⋆^] that are stochastically stable^15^ are exactly those that maximise the potential Φ provided the following conditions apply:

(i) Φ is state invariant
(ii) The action invariant state transition process *P*(*a, ·*) is aperiodic and irreducible over the states *X*(*a*).

Condition (i) is fulfilled since our potential is state-independent by definition.

For all states x and actions a, we have (1) any state can be immediately repeated: *x* ∈ *G*^+^(*a, x*); (2) the set of next states *G*^+^ does not change along an action invariant state trajectory: ∀*x*^*′*^ ∈ *G*^+^(*a, x*) : *G*^+^(*a, x*^*′*^) = *G*^+^(*a, x*); (3) since 𝒢 is connected: |*G*^+^(*a, x*)| > 1.

Condition (ii) is fulfilled since in combination with Eq. 5, (1 ∧ 2) implies that *P* is irreducible and (1 ∧ 3) implies that *P* is aperiodic.

Hence, log-linear learning maximises the pot}ential Φ for *M*.

Log-linear learning is just one form of stochastic hill-climbing in which the individual only assesses its own utility. We will never know the intricate details of the behaviour revision in any natural system in full but convergence in potential games is very robust against changes in the finer details of the behaviour revision protocol [42, 52]. This robustness can give us some confidence that these results carry over to other related forms of biological behaviour adaptation.

## 5 Application to *Eciton* bridge building

To demonstrate that our model, beyond being conceptually plausible, is consistent with real biological scenarios we apply it to the case of Army ant (*Eciton*) bridge building. Army ants form assemblages of their own bodies for a variety of purposes, including to form “bridges” that span gaps in the colony’s foraging trail (Fig. 8). Based on data from field experiments, Reid at al. [53] presented a mathematical model of this behaviour revealing that it is consistent with the assumption that these bridges are built to optimise foraging traffic and transport. The experiments inserted an adjustable angular device (Fig. 9) into natural foraging trails, prompting the animals to bridge across this forced “detour”. Initially, a very small bridge forms at the point where the two arms of the apparatus meet. Subsequently, increasingly many ants join the bridge, allowing it to lengthen and move further down the arms of the apparatus. However, at some point this process stops and the bridge stays approximately in place.

**Figure 8:**
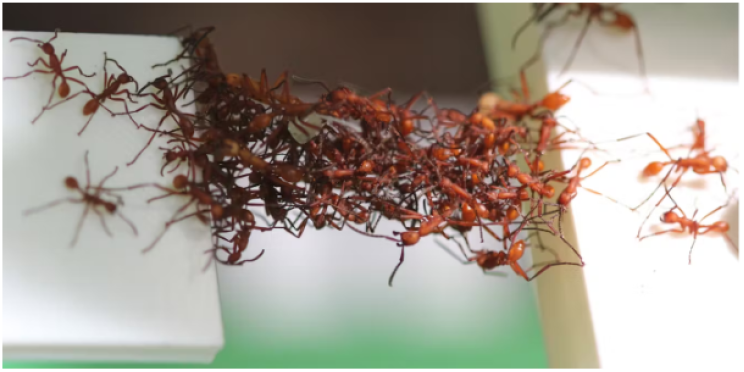
A “Living Bridge” formed by *Eciton* (from [53])

**Figure 9:**
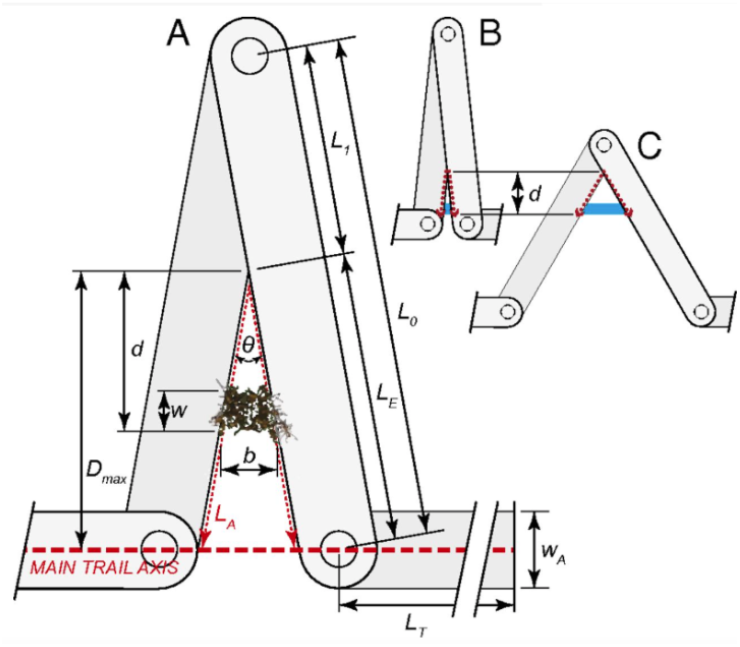
Experimental apparatus (from [53])

The work by Reid at al. explains that the final position of the bridge depends on the angle of the apparatus and is optimal in terms of balancing the benefit of a shorter trail with the cost of sequestering ants from the foraging process to bridge building. While their work elegantly explains why these bridges are built and how they should be positioned, [53] does not offer a mechanistic explanation of how individual workers control their behaviour to achieve this collective benefit, because is not clear that the ants can sense the optimal flow directly.^16^ Here we show that there is no need to make this assumption and that the evolution of objectives can enable the colony to adjust their behaviour for optimal flow even if this cannot be sensed directly.

The hypothesis of [53] is that the ants optimise an (empirical) traffic density that is estimated as

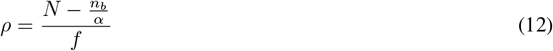

where *N* is the total number of foragers, *n*_*b*_ the number of foragers sequestered into a bridge, and *f* the total travel distance.^17^ *α* is a free parameter that varies between experiments and is supposed to capture various influences on the global density as well as other factors, such as differing body sizes.

From the perspective of an individual worker encountering a bridge, it has to make a decision whether to continue foraging and use the bridge as it finds it or to stop foraging and join the bridge, thereby causing it to be extended.

We show in a simulation that, without sensing *ρ*, objective evolution produces a perception function *N*_θ_ that, when used as the input for this decision making, allows the workers to collectively optimise the transport. Based on earlier research, it is perfectly plausible to assume that a worker on the bridge can sense the local bridge density *s*_1_ = *n*_*b*_/*b*, where *b* is the length of the bridge, as well as the average head-on encounter rate on the trail *s*_2_ = (*N* − *n*_*b*_)/*f*. Our simulation uses these two quantities as the only direct sensory input.

Figure 10 shows the evolved perception functions including the optimal bridge positions (measured in distance from the junction point) according to Reid et al’s mathematical optimisation model (dashed orange vertical line) and as resulting from our simulation model (dashed blue vertical line).

**Figure 10:**
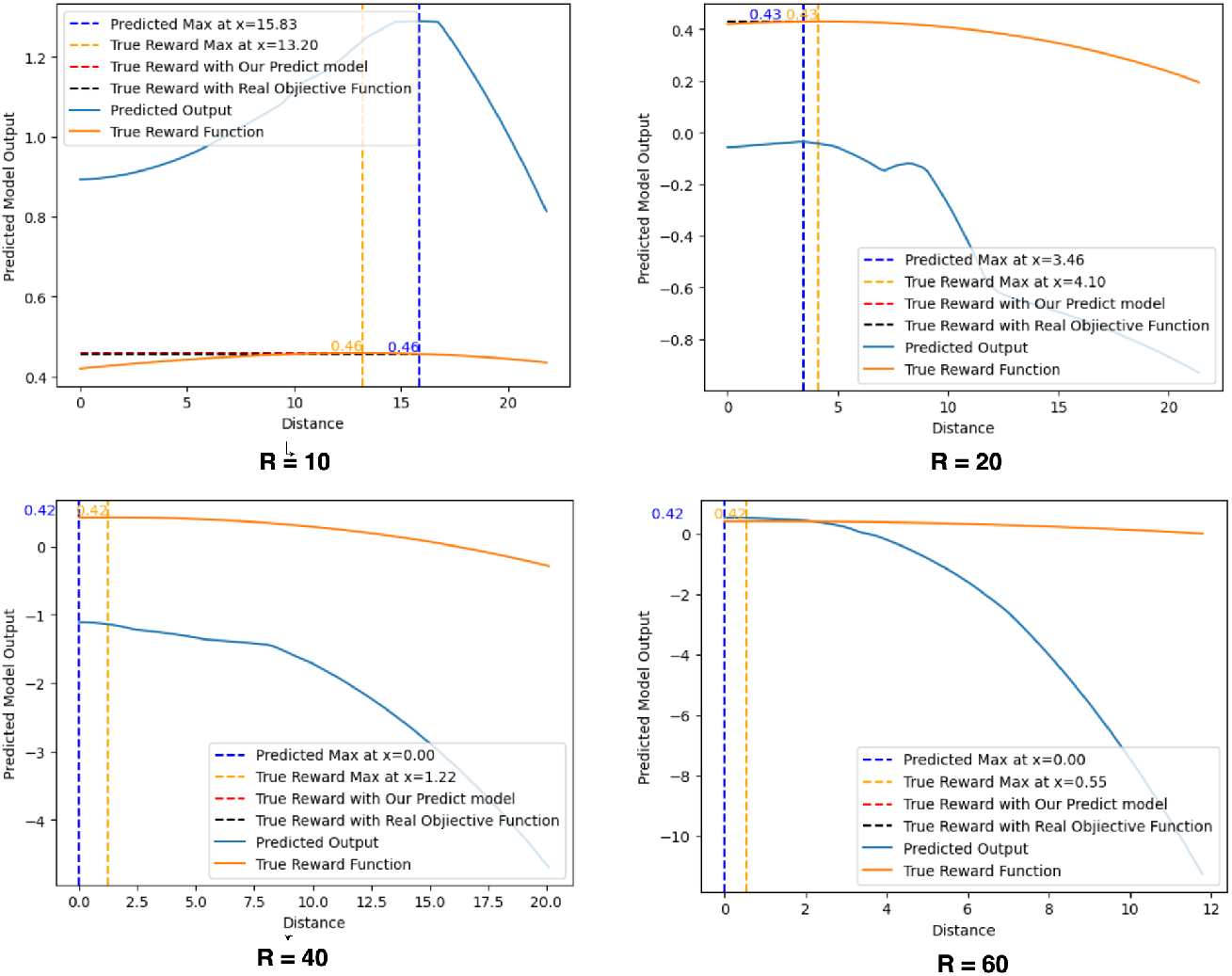
True and evolved perceived objective for *Eciton* bridge building

Figure 11 gives a comparison of the optimal distance in dependence on the angle of the apparatus as predicted by Reid’s optimisation model (orange line) and as reached by objective evolution (blue line). We see that the data is in reasonable agreement.

**Figure 11:**
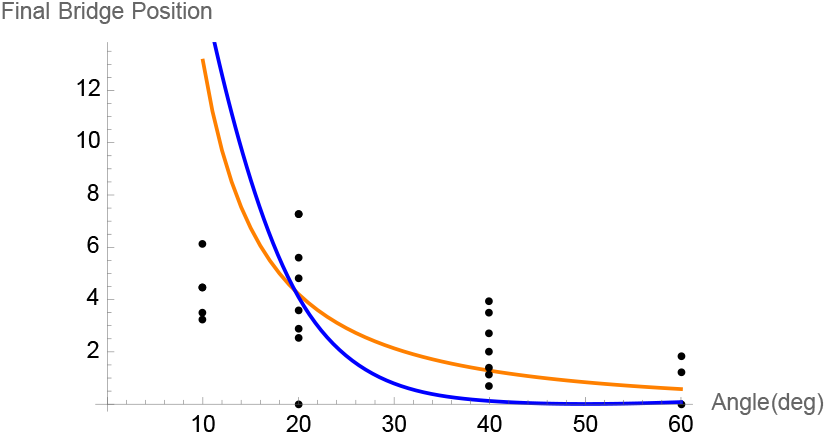
Comparison between the final position of the bridge position predicted by Reid’s model (orange) and our model (blue)in the context of the experimental observations (dots) as a function of the angle *θ* [53].

In summary, we see that objective evolution can offer a possible explanation for the proximate mechanisms individuals use in this concrete biological example to adapt their behaviour so that the collective benefit is optimised even without being able to sense this collective benefit directly and in the absence of any social coordination mechanisms.

## 6 Discussion and conclusions

We have outlined a possible explanation of how pro-social behaviour can be achieved in a collective in the absence of central control by individuals that only make simple decisions based on locally perceived information. Surprisingly, this does not require the individuals to explicitly exchange information regarding the collective benefit. All that is required is the possibility of an evolving perception function that modulates direct sensory input. The only additional assumption that needs to be made for pro-social behaviour to emerge is that the individuals adjust their behaviour via simple hill-climbing mechanisms. The latter is firmly inline with the existing literature.

Interestingly, pro-social decision-making can even be maintained in a complex network of decisions. This is because pro-social atomic decisions allow the relevant information to propagate unhindered throughout the decision network.

We have illustrated how this can work with the example of task allocation, where a colony needs to regulate the allocation of workers to a variety of tasks with competing task demands.

Figure 12 summarises the overall mechanism. Individuals control a decision variable *x* by applying simple learning behaviour according to an evolving perception function *ϕ* and using their directly sensed local information 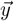. The joint outcome of this regulation determines the global colony fitness *F*. Since the learning mechanisms effectively result in hill-climbing on *ϕ*, the evolutionary pressure causes *ϕ* to assume the shape of a unimodal function that has its peak at the *x* position that optimises the regulation outcome. Since the evolutionary pressure on *ϕ* originates from the collective benefit the maximum of the regulation outcome has to be pro-social. *ϕ* thus establishes an information flow between the individual behaviour and the collective benefit, which cannot be perceived by the individual and is effectively only “visible” at the evolutionary level.

**Figure 12:**
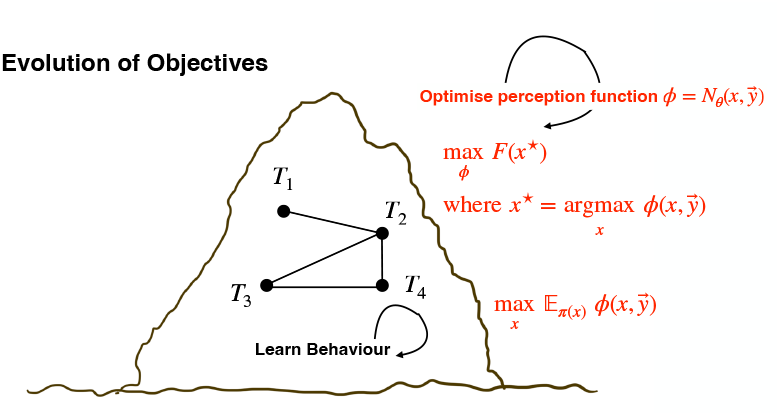
The evolving perception function *ϕ* establishes an information channel from the colony level (fitness) to the decision-making process of the individual.

From the perspective of game theory, *ϕ* is a potential function and reshapes the binary decisions in atomic games as potential games. In this way, it establishes a shared objective so that the pro-social optimum becomes a Nash equilibrium. Since the learning process will reach Nash, rational players, determining their actions according to *ϕ* will behave pro-socially. This reshaping of the individual games turns the whole decision network into a state-based potential game, which enables players to optimise their behaviour in a decentralised fashion.

In a complex task network, rigid better-reply dynamics may only lead to local optimisation, whereas log-linear learning guarantees global optimisation. The core behavioural difference is that log-linear learning allows the agent to occasionally adopt an inferior action, which can be understood as a noisy version of better-reply dynamics. A crucial aspect is that the quality of the steady-state distribution improves consistently with decreasing (but non-zero) noise levels while the time to reach the decision increases. In a strict mathematical sense, perfect optimisation thus requires that the noise level be annealed to zero. However, in a biological system this is arguably not necessary, since not perfect optimisation but only approximate optimisation is required and such “good” behaviour can be achieved without controlling the level of noise exactly.

While our question is closely related to the emergence of altruism, a core question in evolutionary biology, it is subtly different. We are primarily interested in proximate mechanisms, namely how an individual can adjust its behaviour to achieve improved collective benefit if it has no direct information on how its behaviour impacts this. The answers to both questions, however, appear to be intimately connected; maybe not surprisingly so. The missing link in the information flow can only be established by evolution. Arguably the most intriguing aspect is that pro-social behaviour must almost necessarily evolve in a collective in which individuals regulate their behaviour with simple hill-climbing learning under the assumption of an evolving perception function.

There are, of course, many limitations to this model. It certainly does not cover the complexity of real task allocation in social insects in which not all individuals behave identically [31]. We are not attempting to build a concrete qualitative model of task allocation but just to elucidate a new hypothesis of fundamental principles of collective decision making using task allocation as an example, somewhat in the tradition of [37, 54]

Our model rests on the assumption that locally sensed information can be “sufficiently informative” regarding the collective benefit. What exactly “sufficiently informative” means is somewhat open to interpretation. In the strict mathematical optimisation model detailed above we have taken this to mean that the local information can be used to reliably recover the sign of the gradient of the collective benefit in respect to the decision variable. This is a relatively strong assumption but only necessary in this form if we want to show that the true exact optimum will be reached. True optimisation is generally not to be expected and rarely if ever seen in biological behaviour. Thus, it should be possible to relax this assumption and still have a system that develops good albeit not optimal pro-social behaviour based on the same principles.

Could this theory be tested experimentally? It should in principle be possible to pick a species with the potential for fast evolution and adaptation, such as clonal raider ants [55] or potentially Pharaoh ants, and to artificially manipulate the reproductive success of the colony according to controlled task profiles. Modifying nutritional geometry might offer an approach to manipulate task profiles in a meaningful way [56]. In this way it could be shown whether the collective allocation process can be evolutionarily manipulated at the colony level. While this would be extremely interesting, it would also appear to be a formidable experimental undertaking. However, experimental evidence that collective behaviours are shaped by colony-level selection has already been presented for Pharaoh ants [57] and breeding programs, at least in honey bees, are not uncommon [58] and can be successful in relatively few generations [59] so that such experiments do not have to be impossible.

For evaluating this hypothesis in the context of a concrete system, whether ants or any other collectively living organism, a crucial question obviously is whether the required neural circuitry could evolve. While this seems possible in general [60] we are probably quite far from answering this in the affirmative, specifically for ants, since their neuroethology is still in its relatively early stages [61]. Within social insects, it might be more promising to progress such a line of investigation with honey bees, whose brain is much better understood.

Another, not quite as ambitious experimental approach would proceed along the lines of Section 5 by conducting further manipulative experiments in which (a proxy of) collective benefit is clearly quantifiable but the individuals cannot sense this benefit directly. Even though few such experiments seem to exist in the literature, they are clearly feasible as [53] and Burd&Howard’s work on *Atta* [62, 63] have shown. This would provide a more immediate way to gather further experimental support for our hypothesis.

## Acknowledgments

This work was supported in part by ARC Discovery grant DP200100036 to BM. Some parts of this research have previously been published in “Task Allocation and Information Sharing Strategies: A Study of Collective Decision-Making in Ant Colonies”, Yue Yang, PhD thesis Monash University, submitted April 15, 2025.

## Footnotes

1 More correctly, it should be termed “task selection”, since tasks are selected in a self-organised manner by the individuals performing them rather than being “allocated.” However, “task allocation” is the established term.

2 With a few exceptions, such as honeypots [36]

3 Of course, this is not saying that information exchange between individuals could not be used to improve self-organised task allocation.

4 Any other evolvable representation of a universal function approximator would be possible but a neural network seems both a technically sensible and a biologically more relevant choice.

5 For the sake of simpler notation, we will omit the time index in the equations below where it is clear from the context.

6 Note, however, that crowding does no universally decrease foraging efficiency [47].

7 For simplicity, we restrict the action space in such a form that the only tasks accessible to an agent are those in its current game and note that a state-invariant action space could be used instead by appropriately extending the payoff function and transition kernel.

8 Since 𝒢 is undirected, (*T*_*i*_, *T*_*j*_) and (*T*_*j*_, *T*_*i*_) denote the same edge.

9 This assumption is for clarity and notational convenience only and could be relaxed.

10 We acknowledge that this is a relatively small spread of problems. The reason is that the simulation involves the artificial evolution of a large population of agent-based simulations, which is computationally a very costly process.

11 Note that we have trivially overloaded the fitness function *F*at this point, since it was originally defined on an action profile. However, as can easily be seen from Eq. 3, the function essentially works by computing the population fractions from the joint action profile, so the meaning of this should be straightforward. *F*(*β*) = *F*(*Z*) for #_1_(*Z*) = *β*.

12 Strictly speaking, we only need to show that *M*is an *ordinal*state-based potential game for which this can be relaxed to 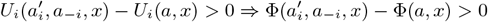.

13 Generally, for state-based potential games the potential is allowed to be state dependant.

14 The theorem actually proves this more generally for a broader subclass of cardinal potential games but we do not require this generalisation here.

15 The set of stochastically stable state of the process is defined as the support of its limiting state distribution for lim_*τ*→0+_.

16 This is mostly because of the free parameter *α*in Eq. 12.

17 We note that for *ρ*to be a density, *f*would have to be an area, not a length. While units would obviously differ, this is numerically equivalent up to a constant under the assumption of a constant trail width. For consistency with [53], we keep *f*a length to enable easier comparison with this work.

